# Isolation, culture and characterization of *Arsenophonus* symbionts from two insect species reveal loss of infectious transmission and extended host range in the apicola-nasoniae group

**DOI:** 10.1101/2022.09.13.507752

**Authors:** Pol Nadal-Jimenez, Steven R. Parratt, Stefanos Siozios, Gregory D.D. Hurst

## Abstract

Vertically transmitted ‘Heritable’ microbial symbionts represent an important component of the biology and ecology of invertebrates. These symbioses evolved from ones where infection/acquisition processes occurred within the environment (horizontal transmission). However, the pattern of evolution that follows transition from horizontal to vertical transmission is commonly obscured by the distant relationship between microbes with differing transmission modes. In contrast, the genus *Arsenophonus* provides an opportunity to investigate these processes with clarity, as it includes members that are obligate vertically transmitted symbionts, facultative vertically transmitted symbionts, strains with mixed modes of transmission and ones that are purely horizontally transmitted. Significantly, some of the strains are culturable and amenable to genetic analysis. We first report the isolation of *Arsenophonus nasoniae* strain Pv into culture from the ectoparasitic wasp *Pachycrepoideus vindemmiae* and characterize the symbiosis. We demonstrate maternal vertical transmission and find no evidence for paternal inheritance, infectious transmission or reproductive parasitism phenotypes. This leads us to conclude this strain, in contrast to related strains, is a facultative heritable symbiont which is likely to be beneficial. We then report the serendipitous discovery and onward culture of a strain of *Arsenophonus* (strain Pb) from the blue butterfly, *Polyommatus bellargus*. This association extends the range of host species carrying *A. nasoniae/A. apicola* symbionts beyond the Hymenoptera for the first time. We perform basic metabolic analysis of the isolated strains using Biolog plates. This analysis indicates all strains utilize a restricted range of carbon sources, but these restrictions are particularly pronounced in the *A. nasoniae* Pv strain that is solely vertically transmitted. Finally, we demonstrate the *Arsenophonus* strain Pb from the blue butterfly can infect *Galleria* waxworms, providing a model system for investigating the functional genetics of *Arsenophonus-insect* interactions.

## INTRODUCTION

Heritable symbionts – microbes that are vertically transmitted from parent to offspring – represent an important component of the biology of invertebrates (1). These symbionts can provide services to the host in the form of essential nutrient supplementation (2), natural enemy protection (3), and thermal adaptation (4). Further, their capacity to block viral replication (5, 6) enable their use in pest and vector control, as a public health intervention interrupting the competence of arbovirus vectors (7, 8). In some cases, they are obligately required by the host for function, and in most cases the microbes are fastidious and unable to grow outside of a host environment. Contrastingly, heritable microbes can also act as reproductive parasites, causing sex-ratio distortion and cytoplasmic incompatibility (9). The impact symbionts have on individual hosts – both beneficial and parasitic - cascades to ecological and evolutionary dynamics, for instance driving changes in the natural enemy-host dynamics (10), altering patterns of sexual selection (11), and potentiating host biodiversity over macroevolutionary timescales (12).

Vertically transmitted symbioses originally evolved from symbioses involving horizontal (infectious) transmission (13), which either involved infection of a host by an environmental microbe, host capture of an environmental microbe, or an interplay between the parties (1). Vertical transmission from parent to offspring would then have evolved as a novel means by which hosts retain useful symbionts, or as a mechanism for microbes to acquire new hosts (13). Investigating the processes that accompany transitions in symbiont transmission mode remains a key challenge, as existing vertically transmitted microbes are commonly evolutionary distant from horizontally transmitted species, obscuring the processes that occur early in the evolution of vertical transmission. In order to understand the processes that occur on first evolution into vertical transmission, we must compare symbionts with vertical transmission to closely related to strains where symbiosis establishes horizontally through the environment. The contrasts then allow us to establish the tempo and mode of gene loss during vertically transmitted symbiosis, as well as allowing comparative analysis of the properties of different modes of symbiosis.

*Arsenophonus* is a microbial genus engaged in symbiotic interactions with diverse arthropod host species. Within the clade there are horizontally transmitted species (*A. apicola*) (14, 15), symbioses with mixed modes of transmission (*A. nasoniae* and insect-vectored *Cand*. A. phytopathogenicus plant pathogens) (16, 17), vertically transmitted symbionts that are not required for host function (e.g. *Cand*. A. triatominarum) (18), and obligately required vertically transmitted symbionts (e.g. *Cand*. A. arthropodicus and *Cand*. A. lipopteni from hippoboscid flies) (19, 20). The relationships vary from likely pathogenic (*A. apicola* in honey bees), through reproductive parasitism (*A. nasoniae* in *Nasonia* wasps), to mutualism (*Cand*. A. arthropodicus and *Cand*. A. lipopteni from hippoboscid flies). A key feature enabling functional analysis in this clade is the ability to isolate and grow some of the strains within this clade in cell-free culture, with *A. nasoniae* and *A. apicola* both amenable to culture and genetic manipulation (15, 21). *Arsenophonus nasoniae* itself also represents a fascinating microbe, possessing a very complex genome containing a high density of prophage elements and hyperdiverse extrachromosomal elements (22).

In this paper, we report the isolation and characterization of two strains of *Arsenophonus* from the *A. nasoniae/A. apicola* subclade that to date is dominated by strains which retain at least some infectious transmission. Previous work had reported *Arsenophonus* infection in *Pachycrepoideus vindemmiae*, an ectoparasitic wasp of fly pupae (23); we isolated this infection to pure culture and characterized its phenotype in the wasp host in terms of transmission and reproductive parasitism. We then report the serendipitous isolation of an *A. nasoniae/A. apicola* relative from the adonis blue butterfly, *Polyommatus bellargus*. We then compare their carbon utilization sources and inhibitory factors for these strains, using Biolog plates to investigate if transition to strains with pure vertical transmission have reduced metabolic flexibility. Finally, we report their capacity to grow in *Galleria melonella* waxworms, a model system for understanding the mechanistic basis of host-microbe interactions.

## MATERIALS AND METHODS

### Isolation and characterization of *Arsenophonus* spp. in *Pachycrepoideus vindemmiae*

The *P. vindemmiae* line used in these experiments was derived from a single *Arsenophonus-infected* female caught near Pierrefeu, south East France by F.Vavre *et al*, University of Lyon, France (23). *Pacycrepoideus vindemmiae* is a *Drosophila* pupal parasitoid that lays a single egg in a fly host. Isolation of the symbiont was achieved by extracting wasp pupae from their drosophilid hosts, surface sterilizing them with 70% EtOH, washing with ddH_2_O and homogenizing in sterile PBS. Homogenate was then spread onto cell-free media as described previously (24) and allowed to grow at 25°C for 4-6 days until single colonies were visible. A single clone was then picked into 100 μl of sterile PBS, 20 μl aliquots of which were spread onto fresh GC agar plates using sterile resin beads to form bacterial lawns after a second bout of incubation and growth. Microbe identity was confirmed through colony PCR amplification of the 16S rRNA amplicon using primers 27F and U1492R (25), sequencing the amplicon and performing a nucleotide BLAST search with the resulting sequences. The resultant strain was named *A. nasoniae* strain Pv.

We investigated the transmission biology of *Arsenophonus* in *P. vindemmiae* and tested for the presence of reproductive parasitism, investigating a) the efficiency of vertical transmission b) the capacity for infection to spread between lineages through superparasitism c) the presence of maternal vs paternal inheritance, sex ratio distortion and incompatibility phenotypes.

To estimate the vertical transmission efficiency of *A.nasoniae* Pv, infected female *P. vindemmiae* wasps were collected as pupae from stock vials, allowed to eclose and mate for 24 hours with 3 infected males from their natal line. These females were then given *ad libitum* (100+) *D. melanogaster* (Canton S) pupae in which to oviposit individually for 48 hours. After this time females were collected and their infection verified by PCR screening specific to *A. nasoniae* based on the 16S rRNA gene using the primers Arse16S-F and Arse16S-R (26).

The adult progeny of infected females were collected after 30 days, allowing for full development and eclosion. Fifteen female progeny from each mother were then selected at random for PCR screening for infection to generate a transmission efficiency value per mother. DNA quality was verified for each sample by amplifying a portion of the insect mitochondrial COI gene (Primers: **LCO**. 5’ GGT CAA CAA ATC ATA AAG ATA TTG G 3, **HCO**. 5’ TAA ACT TCA GGG TGA CCA AAA AAT CA 3’) (27). Five samples failed to amplify a product for these control primers to a visible standard through gel electrophoresis, so were discarded from analysis.

Previous work has established horizontal transmission of *A. nasoniae* in another wasp, *N. vitripennis* when both infected and uninfected mothers share a host pupa. In *N. vitripennis*, vertical transmission is inefficient, and horizontal transmission is necessary for maintaining the symbiont infection (17). To test whether *A. nasoniae* Pv also transmits horizontally between lineages of *P. vindemmiae*, we cohoused *Arsenophonus-infected* wasps with isogenic antibiotic-cured uninfected wasp females (line established > 5 generations previously) and scored infection prevalence in the emerging offspring. To this end we followed a similar procedure as above, but allowed four female wasps to simultaneously co-lay in the same vial of *Drosophila* pupae at a 2:2, *Arsenophonus* positive (*A+): Arsenophonus* negative ratio (*A-*). Once females were grouped it was no longer possible to distinguish *A+* from *A*-. Therefore, following oviposition for 48 hours, all mothers were collected and screened for infection (80 total) resulting in three being discarded because their observed infection ratio in mothers was lower than the expected 2:2, giving a total of 17 replicates for analysis. We collected adult progeny that emerged from these cohoused lays after 30 days and PCR tested a random sample of 30 female offspring for *Arsenophonus*. Our null expectation under the assumption that Pv only transmits vertically is that 50% of offspring emerging from each replicate will be infected with *Arsenophonus*.

*A. nasoniae* is as a sex-ratio distorter in the parasitoid *Nasonia vitripennis*, whilst other heritable symbionts distort sex ratios by inducing cytoplasmic incompatibility. To test for sex ratio distortion and cytoplasmic incompatibility in *P. vindemmiae* – Pv infections, we performed a set of 2 × 2 factorial crosses of infected and uninfected male and female wasps. These crosses allow us to test for:

i. Paternal and maternal transmission efficiency of *Arsenophonus*, which would be shown by *Arsenophonus-infection* in progeny of *A-* female x *A+* male crosses and A+ female x A-male crosses respectively.
ii. Any sex ratio distortion phenotypes, which would be shown by differences in offpsring sex ratios produced by *A+* females compared to *A-*.
iii. Any cytoplasmic incompatibility, which would be shown if progeny from *A-* mothers crossed to *A+* fathers are either all male or have higher rates of failure to emerge as adults.

To test these hypotheses, males and females for the crosses were collected within 24 hours of eclosion and kept in mating groups of 3 males and 3 females according to treatment, with access to honey water for nutrition. We allowed mating in groups to reduce the impact any infertile or unresponsive males in the experiment. After mating, females were removed and placed into individual ‘host vials’ with ten freshly pupated *D. melanogaster* (CS) (11 days old at 25°C) on removable plastic sticks embedded in their culture vials. Infection status of the parents was confirmed *post hoc* by PCR screening as described above. Final sample sizes of each cross after removal of replicates that died during oviposition or were not of the assigned infection status were as follows: 11 x both parents infected, 23x infected fathers only, 16 x infected mothers only, 12 x neither parent infected. The number and sex ratio of the F1 offspring were scored for each clutch as they emerged. Where possible, up to two live F1 female offspring from each brood were taken for PCR screening for *A. nasoniae*. This was to determine whether transmission of the infection was maternal, paternal or biparental. The total number of offspring screened per treatment were: N=12 both parents infected, N=24 infected fathers, N=16 infected mothers, N=10 neither parent infected.

### Data Analyses

Infection prevalence under vertical transmission is reported as the observed proportion of offspring that tested positive for the symbiont, including 95% confidence intervals.

To test for horizontal transmission when A+ and A-wasp females share a group of fly hosts we used a one-sample Z-test to compare the total proportion of emerging offspring that were infected with the symbiont against the null expectation of 50% infection prevalence.

To test for differences in infection prevalence, sex ratio, and offspring viability between broods produced by crosses of differentially infected parents we used generalised linear models with binomial errors. In all cases the infection status of the parents was fit as categorical independent variable and the respective response variable as a binomial dependant variable. For infection prevalence, due to screening a maximum of two offspring per clutch, we grouped all individuals by treatment level prior to analysis. For brood viability and sex ratio we included a random intercept in our models to account for the nested nature of broods within treatments. Models were fit with ‘lme4’ in R and significance levels of main effects generated with type II Wald Chi square tests implemented by ‘car::Anova()’. All probabilities and 95% confidence intervals were calculated with ‘binom::binom.confint()’ using the logit method.

### Serendipitous isolation of *Arsenophonus* infection from the butterfly *Polyommatus bellargus*

Illumina reads derived from thirty male *P. bellargus* collected in August 2017 from Swellshill, Gloucestershire, UK were checked for the presence of microbes using Phyloflash (28). The reads from one specimen that died shortly after collection indicated the presence of *Arsenophonus* reads at very high ratio (~11.5%), and remaining abdomen material for this specimen was retrieved from the −80°C freezer. Material was reconstituted in BHI. Identity of the colony as *Arsenophonus* was confirmed through PCR assays combined with sequencing of the 16S rRNA amplicons, as described above.

### Relatedness of strains

We estimated the relatedness of strains using the sequence of three marker genes: *fbaA, yaeT* and *ftsK* previously used to type *Arsenophonus* strains (23). Marker sequences for *Arsenophonus* strain Pb were amplified and Sanger sequenced using the primers from Duron et al. (23). Sequences were aligned against those of existing strains at protein level and then back-translated to nucleotide. Alignment columns with more than 40% gaps were removed. Best fitting model estimations was performed using ModelFinder (29) as implemented in IQTREE (30). Phylogenetic relatedness of strains was estimated through Maximum Likelihood method, using IQTREE and the TN+F+G4 model with 1000 ultrafast bootstrap replicates (31).

### *In vitro* growth requirements of strains isolated

BIOLOG GEN III plates (Cat. No. 1030) were used to ascertain the *in vitro* physiological and biochemical characteristics of *Arsenophonus nasoniae* strain Pv*, Arsenophonus* strain Pb, with comparison to *A. nasoniae* Nv and *A. apicola*. For the BIOLOG GEN III assays, we used IF-A inoculating fluid (Biolog, Cat. No. 72401). All *Arsenophonus* spp. were grown for four to six days (till a maximum OD600 = 0.4-0.6) in brain heart infusion (BHI) broth (Oxoid, UK) at 30°C and 250 rpm and BIOLOG GEN III plate set up as previously described (15). The plate was subsequently incubated at 30°C without shaking during six to eight days before scoring according to manufacturers instructions.

### Capacity to grow in *Galleria*

*Arsenophonus nasoniae* strain Pv and *Arsenophonus* strain Pb were independently transformed with plasmid *pOM1::gfp* using previous protocols to establish GFP expressing clones. The capacity of these strains to infect was examined alongside GFP expressing *A. nasoniae* (21) and a GFP-expressing *E. coli* MG1655 harbouring the same plasmid. In brief, strains were grown in BHI agar plates for 6 days (Nv and Pv); 2 days (Pb) and overnight (*E. coli* MG1655), and then subcultured in BHI broth for 2 (Nv, Pv) and 1 days (Pb and *E. coli*). 1 μl of culture (OD_600_=0.4-0.6) was injected into 25 individual second instar *G. mellonella* larvae using a nanoject III and remains of culture retained for estimation of bacterial density through serial dilution; in all cases, >10^5^ cfu were introduced. Sham (BHI) infected negative controls were also completed. Individual *G. mellonella* larvae were then monitored for visible infection, scored as expression of GFP, over 3 days at x 25 °C.

## RESULTS

### Isolation and characterization of *Arsenophonus* nasoniae from *Pachycrepoideus vindemmiae*

*Arsenophonus nasoniae* was successfully isolated from *P. vindemmiae* to pure culture. Colonies were slow growing (5 days at 30 °C), and morphology resembled *A. nasoniae*. 16S rRNA sequence was obtained (Accession number OP289003) and confirmed the isolates as closely related to *A. nasoniae*, and the strain named *A. nasoniae* strain Pv.

The *Arsenophonus nasoniae* strain Pv in its native host *P. vindemmiae* exhibited high vertical transmission efficiency and no evidence of significant levels of horizontal transmission (Figure 1A, 1B). When only a single infected female produces offspring, vertical transmission efficiency of *Arsenophonus* is estimated at 98.3% (Figure 1A). When two infected and two uninfected females share access to the same group of fly pupal hosts, the mean infection prevalence of wasp offspring emerging from the group is 53.3%. This is not a significant deviation from the 50:50 infected/uninfected ratio we would expect to see for vertical transmission where half the female parents are infected (P = 0.453, one-sample Z-test of proportions, confidence intervals: 0.465 - 0.581).

**Figure 1:**
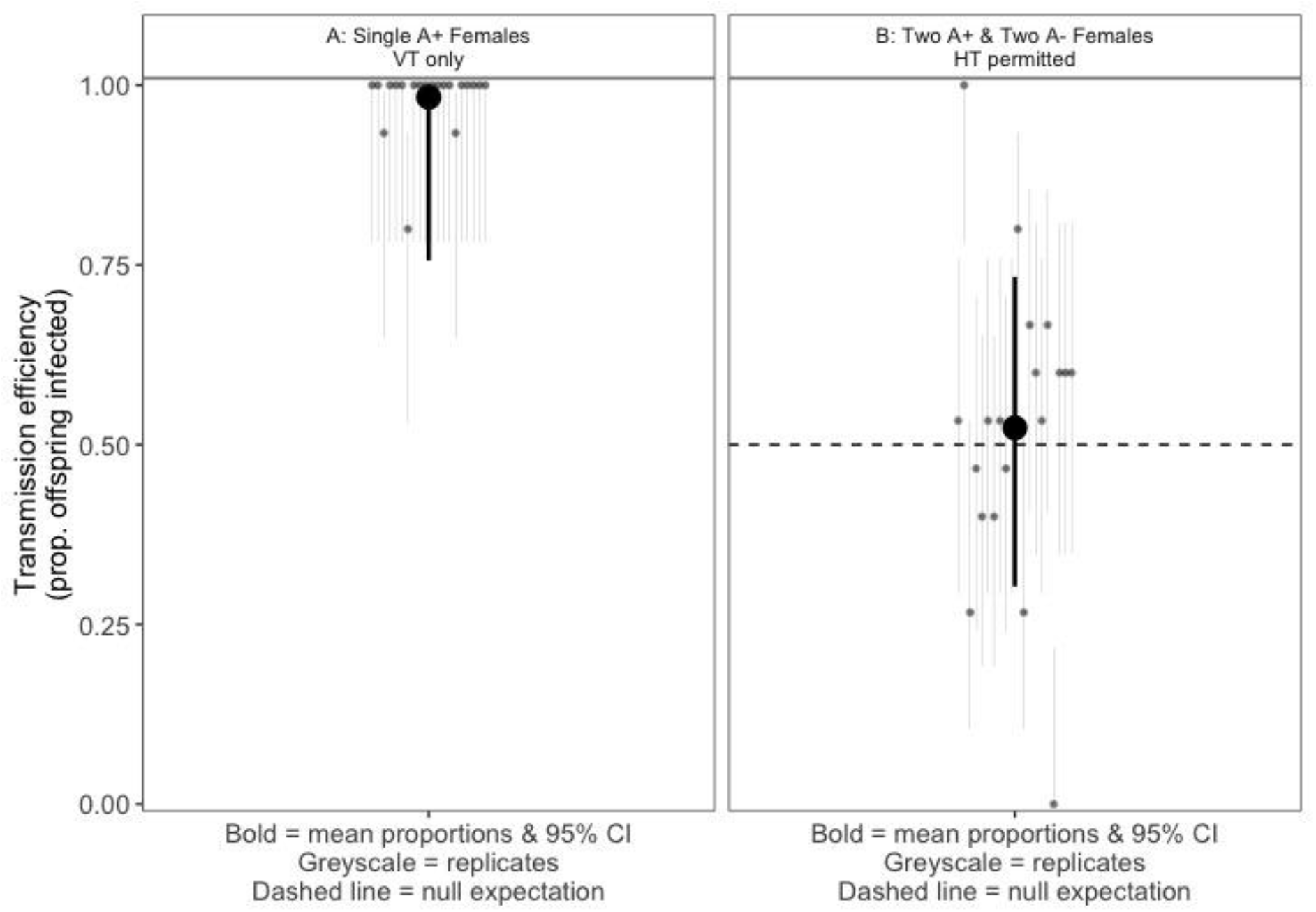
A) Vertical transmission of *Arsenophonus nasoniae* in *Pachycrepoideus vindemmiae* has a high mean efficiency of 98.3%. Light grey points and errors represent infection prevalence of 20 replicate broods laid by lone infected mothers (15 randomly selected female offspring screened from each brood). Dark point and errors show the mean proportion of progeny infected and mean 95% confidence interval calculated using the logit method. B) There is no evidence of significant horizontal transmission, or a (dis)advantage to infection by *Arsenophonus nasoniae* in *Pachycrepoideus vindemmiae*. Pale points and errors represent infection prevalence in 20 replicate broods co-laid by 2:2 infected:unifected mothers (15 randomly selected female offspring screened from each brood. Dark points and errors represent mean infection prevalence and upper and lower 95% confidence intervals calculated with the logit method. Dashed line illustrates the null expectation of 50% infection if there is no HT and no (dis)advantage to symbiont infection.

Vertical transmission was solely maternal. Infected progeny were observed only when the female parent was infected, with none of 24 offspring infected where the father was infected and the mother was not (95% Binomial Confidence intervals on paternal transmission rate: lower = 0.0, upper = 0.142)(Figure 2A). Complete separation of infection across treatment (i.e. no infection observed in broods with infected fathers only or in uninfected control broods) prevented statistical test of this differences. We also found evidence for heterogeneity in clutch sex ratio between any of the treatments (Wald χ^2^ = 2.1, df = 3, *P* = 0.55, mixed effect logistic regression model with replicates fitted as random intercepts)(Figure 2B). This is strong evidence against *A. nasoniae* causing CI or other sex ratio distortion phenotypes such as feminization or male-killing. Under CI, male-biased sex ratios are expected where the male partner was infected and the female partner uninfected (as male progeny in Hymenoptera are not the product of sex they are not affected by CI) (32). Other sex ratio distorting phenotypes should produce female-biased sex ratios when the female was infected irrespective of paternal infection status.

**Figure 2:**
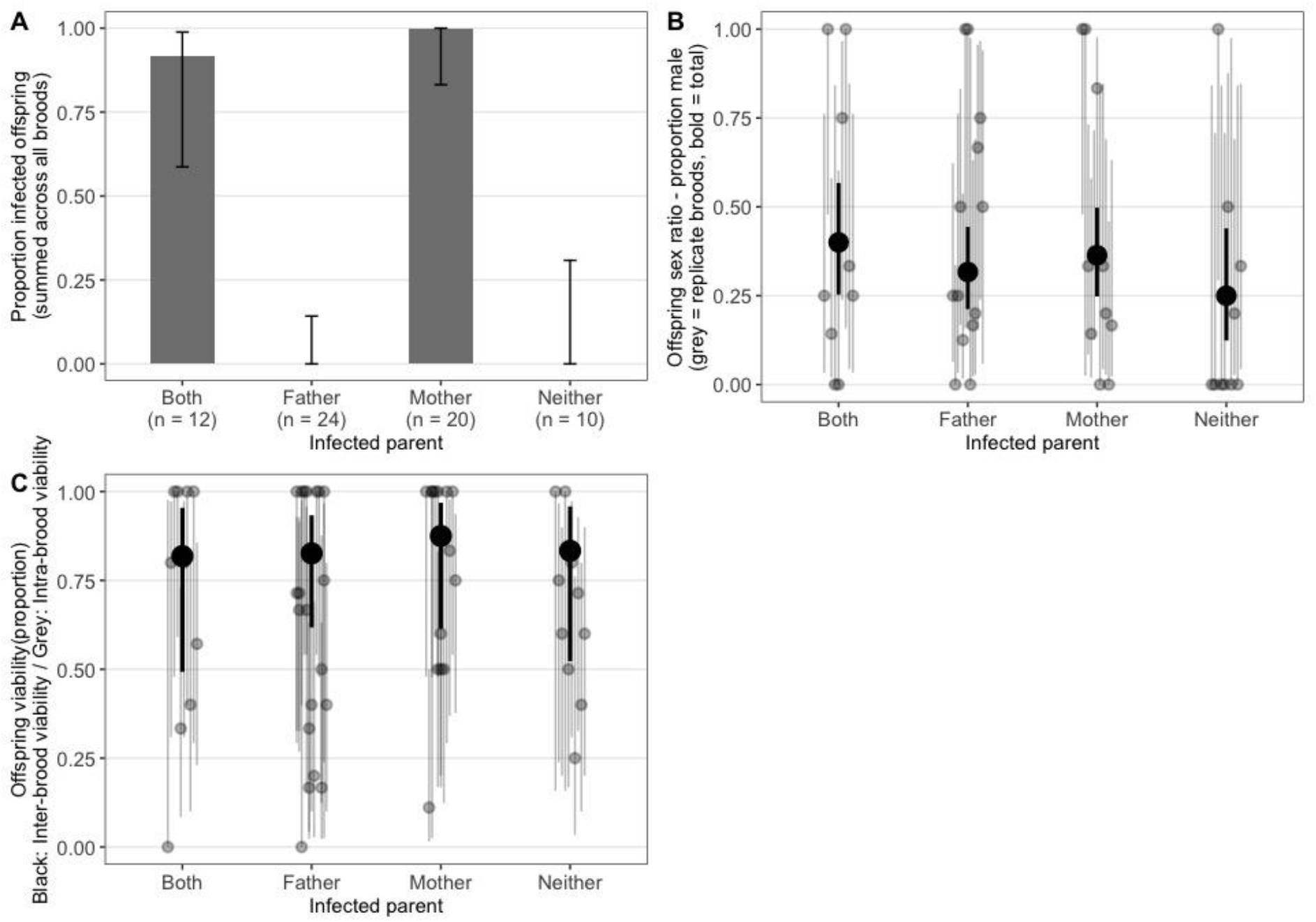
A) Proportion of *Arsenophonus* positive progeny from different combinations of parental infection stats: both mother and father infected, father only, mother only, and neither. Results are given as the mean of replicate crosses, with error bars representing 95% binomial confidence intervals. B) Offspring sex ratio (proportion male) from crosses with different infection status. Grey dots indicate individual families, Black dot represents mean across replicate families. C) Offspring viability (proportion of pupae successfully parasitised with emerging wasp). Grey dots indicate individual families, Black dot represents mean across replicate families.

Finally, we also find no significant differences in clutch viability between treatments, (Wald χ^2^ = 0.23, df = 3, *P* = 0.97, Figure 2C) which further indicates that infection with *Arsenophonus* does not cause incompatibilities between differentially infected parents nor extreme pathogenicity in the wasp.

### Isolation of *Arsenophonus* spp. from the butterfly *Polyommatus bellargus*

*Arsenophonus* isolation was obtained from −80 °C frozen material from the butterfly *Polyommatus bellargus* and growth in liquid culture was achieved easily (henceforth, *Arsenophonus* strain Pb). The strain was closely allied to *A. nasoniae/A. apicola* based on 16S rRNA sequence (OP203938). The relatedness of this strain to others was estimated based on concatenated *fbaA, ftsK* and *yaeT*. This analysis indicated with high certainty that the Pb strain lay within the nasoniae-apicola clade in which other culturable *Arsenophonus* lie. The strain is sister to other known *A. nasoniae* strains (Figure 3).

**Figure 3:**
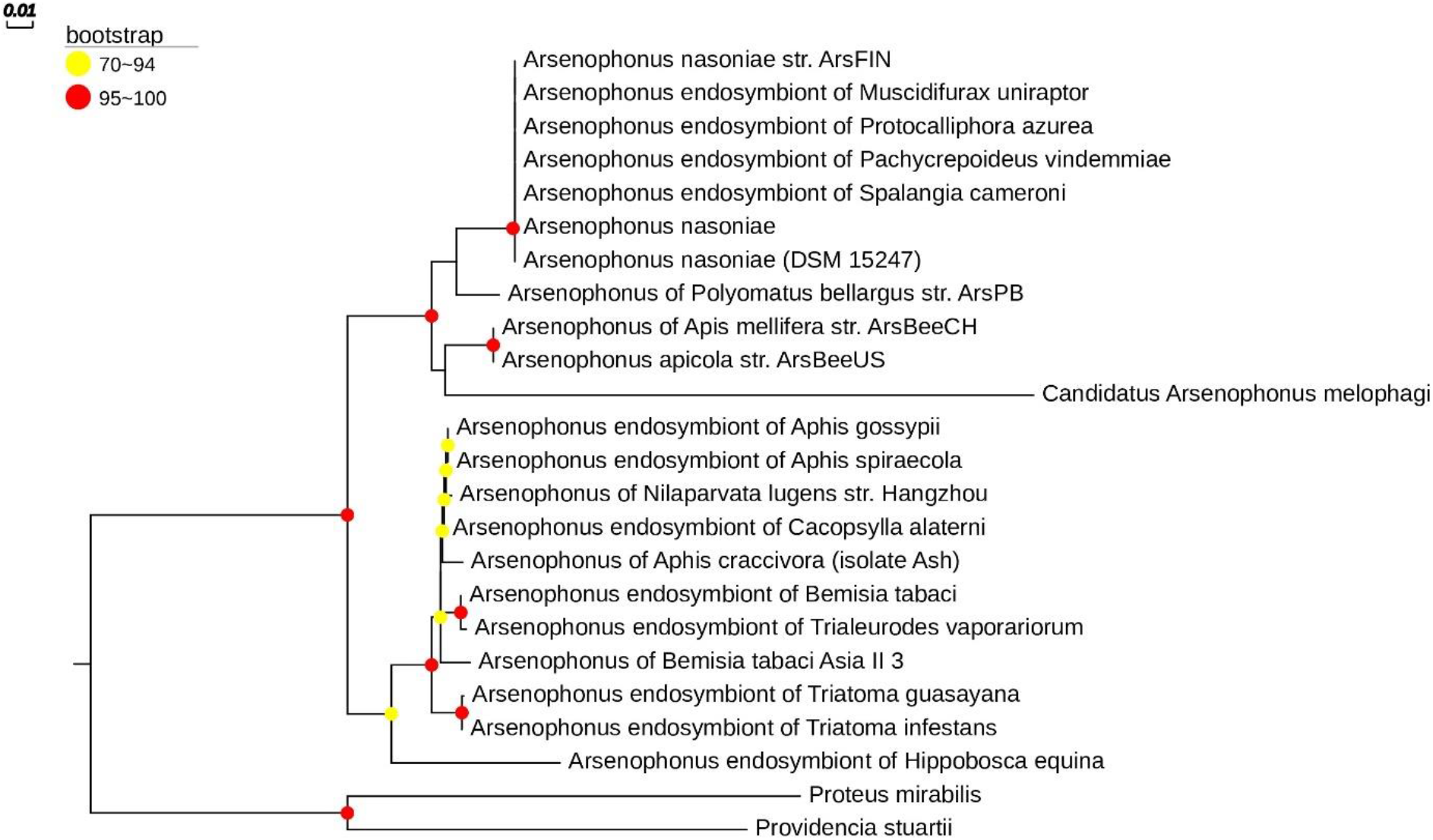
Estimation of *Arsenophonus* phylogeny using Maximum likelihood (ML) based on the concatenated sequences of *fbaA, ftsK* and *yaeT* genes (as in Duron et al. 2010). Only bootstrap values higher than 70% are shown. The phylogenetic tree was constructed using IQTREE and the TN+F+G4 model. Accession numbers on which the phylogeny is based can be downloaded at https://doi.org/10.6084/m9.figshare.20628159.v1.

### *In vitro* growth requirements of strains isolated

We compared carbon utilization and inhibition across *Arsenophonus nasoniae* isolated from UK *N. vitripennis* (*A. nasoniae* Nv, mixed modes of transmission), from *P. vindemmiae* (*A. nasoniae* Pv, vertical transmission) and *Arsenophonus* Pb from *P. bellargus*, with previously established growth conditions for *A. apicola* (horizontal transmission only). All strains utilized a narrower breadth of carbon sources than *A. apicola*, with the Pv strain for *P. vindemmiae* having the narrowest range (Table 1). *Arsenophonus nasoniae* strain Nv from *Nasonia vitripennis* had the broadest capacity to thrive in the presence of inhibitory compounds and the Pv strain was most affected by inhibitory conditions and compounds (Table 2).

**Table 1:** Carbon utilization sources for diverse *Arsenophonus* strains as estimated from biolog plate analysis: *A. nasoniae* Nv, strain Pv from *P. vindemmiae*, strain Pb from *P. bellargus*. Data from *A. apicola* are given for comparison. The following carbon sources were not utilized by any of the strains: Dextrin, D-maltose, D-Trehalose, D-Cellobiose, Gentiobiose, Sucrose, D-Turanose, Stachyose, D-raffinose, α-D-lactose, D-melibiose, B-Methyl-D-Glucoside, D-Salicin, *N*-Acetyl-B-D-Mannosamine, N-Acetyl-D-Galactosamine, N-Acetyl Neuraminic Acid, D-Galactose, 3-Methyl Glucose, D-Fucose, L-Fucose, L-Rhamnose, Inosine, D-sorbitol, D-mannitol, D-Arabitol, myo-inositol, Glycerol, D-Aspartic Acid, D-Serine, gelatin, L-Alanine, L-Arginine. L-Histidine, L-Pyroglutamic Acid, Pectin, D-Galacturonic acid, L-Galactonic Acid Lactone, D-Glucuronic Acid, Mucic Acid, Quinic Acid, D-Saccharic Acid, ρ-hydroxy phenyl acetic acid, D-Lactic Acid Methyl Ester, citric Acid, Tween-40, gamma-Amino-Butryric acid, B-Hydroxy-D-L-Butyric Acid, Propionic Acid, Formic Acid.

|  | *A. nasoniae*<br>strain Nv | *A. nasoniae* strain<br>Pv | *Arsenophonus*<br>strain Pb | *Arsenophonus*<br>*apicola* |
| --- | --- | --- | --- | --- |
| *D-glucose* | +++ | +++ | +++ | +++ |
| *D-mannose* | +++ | - | +++ | +++ |
| *D-fructose* | +++ | +++ | +++ | +++ |
| *N-acetyl glucosamine* | +++ | +++ | +++ | +++ |
| *D-glucose-6-PO<sub>4</sub>* | +++ | +++ | - | +++ |
| *D-fructose-6-PO<sub>4</sub>* | +++ | +++ | - | +++ |
| *L-malic acid* | +++ | +++ | - | +++ |
| *D-malic acid* | - | - | - | +++ |
| *bromo-succinic acid* | +++ | ++ | - | +++ |
| *D-gluconic acid* | - | - | +++ | +++ |
| *Glucoronamide* | - | - | - | ++ |
| *methylpyruvate* | ++ | - | +++ | +++ |
| *L-aspartic acid* | - | - | +++ | +++ |
| *L-glutamic acid* | ++ | - | +++ | ++ |
| *α-keto-glutaric-acid* | ++ | - | - | +++ |
| *α-keto-butyric-acid* | ++ | - | - | ++ |
| *α-hydroxy-butyric-acid* | - | - | - | ++ |
| *Acetic acid* | - | - | +++ | - |
| *L-serine* | - | - | +++ | - |
| *glycyl-L-proline* | - | - | ++ | - |
| *Acetoacetic acid* | - | - | - | ++ |

**Table 2:** Impact of environmental and xenobiotic stress conditions on growth of *Arsenophonus* strains on Biolog III plates. +++ = maintains full growth under condition stated - = no growth under condition stated.

|  | *A. nasoniae* strain Nv | *A. nasoniae* strain Pv | *Arsenophonus* strain Pb | *Arsenophonus apicola* |
| --- | --- | --- | --- | --- |
| *pH 6* | +++ | ++ | +++ | +++ |
| *pH 5* | ++ | - | - | ++ |
| *1% NaCl* | +++ | ++ | +++ | +++ |
| *4% NaCl* | +++ | ++ | - | +++ |
| *8% NaCl* | +++ | ++ | - | +++ |
| *1% Sodium lactate* | +++ | ++ | +++ | +++ |
| *Fusidic acid* | +++ | ++ | +++ | +++ |
| *D-Serine* | +++ | ++ | +++ | +++ |
| *Troleandomycin* | +++ | ++ | +++ | +++ |
| *Rifamycin SV* | +++ | ++ | +++ | +++ |
| *Minocycline* | +++ | ++ | +++ | +++ |
| *Lincomycin* | +++ | ++ | +++ | +++ |
| *Guanidine HCl* | +++ | ++ | +++ | +++ |
| *Niaproof 4* | ++ | - | +++ | - |
| *Vancomycin* | +++ | ++ | +++ | +++ |
| *Tetrazolium Violet* | ++ | ++ | +++ | - |
| *Tetrazolium Blue* | +++ | ++ | +++ | +++ |
| *Nalidixic Acid* | +++ | ++ | +++ | +++ |
| *Lithium Chloride* | +++ | ++ | +++ | +++ |
| *Potassium Tellurite* | +++ | ++ | +++ | +++ |
| *Aztreonam* | +++ | ++ | +++ | +++ |
| *Sodium Butyrate* | +++ | ++ | +++ | +++ |
| *Sodium Bromate* | - | - | - | - |

### Capacity to grow in a *Galleria* infection model

Neither *A. nasoniae* strain Nv, nor *A. nasoniae* strain Pv, established infection in the *Galleria* model, as measured through GFP fluorescence of injected larvae 3 days post inoculation. In contrast, *Arsenophonus* strain Pb was capable of propagating in *G. mellonella*, with 22 of 25 injected individuals developing a disseminated infection (Table 3). *Galleria mellonella* mortality was observed solely in 5 out of 25 larvae infected with the *E. coli* MG1655 control, and this was in the first 24 h post injection. No mortality was observed over the 72h period in any of experimental treatments.

**Table 3:** Alive/dead status, and evidence of proliferation of microbes from GFP fluorescence, in *G. mellonella* larvae injected with various strains of *Arsenophonus*, assessed daily over 3 days. *E. coli* MG1655 not capable of proliferation, and BHI medium injection are given as controls. N=25 for all cases except BHI control, N=20.

|  | Day 1 |  |  | Day 2 |  |  | Day 3 |  |  |
| --- | --- | --- | --- | --- | --- | --- | --- | --- | --- |
|  | Alive<br>GFP<br>+ve | Alive<br>GFP -ve | Dead | Alive<br>GF +ve | Alive<br>GFP -ve | Dead | Alive<br>GF +ve | Alive<br>GFP -ve | Dead |
| *A. nasoniae* str. Nv | 0 | 25 | 0 | 0 | 25 | 0 | 0 | 25 | 0 |
| *A. nasoniae* str. Pv | 0 | 25 | 0 | 0 | 25 | 0 | 0 | 25 | 0 |
| *Arsenophonus* str. Pb | 20 | 5 | 0 | 23 | 2 | 0 | 23 | 2 | 0 |
| *E. coli* MG1655 | 0 | 20 | 5 | 0 | 20 | 5 | 0 | 20 | 5 |
| BHI control | 0 | 20 | 0 | 0 | 20 | 0 | 0 | 20 | 0 |

## DISCUSSION

The genus *Arsenophonus* comprises members that have horizontal (infectious) transmission (e.g. *A. apicola* in *Apis mellifera* honeybees) (14), reproductive son-killer parasites with mixed modes of transmission (*A. nasoniae* in *N. vitripennis* wasps) (17), and a range of facultative and obligate vertically transmitted symbionts. In this paper, we examined the biology of two symbionts in the *apicola-nasoniae* group. We first ascertained the nature of the symbiosis between *A. nasoniae* Pv and the ectoparasitic wasp *P. vindemmiae*. We then cultured *Arsenophonus* from the butterfly *P. bellargus*, in this case a serendipitous isolation from a −80 °C preserved butterfly abdomen. Finally, we compared the breadth of carbon utilization sources of the isolates to *A. nasoniae* and *A. apicola*, and capacity to infect *G. mellonella* waxworms.

*Arsenophonus nasoniae* strain Pv showed high (98%) maternal transmission fidelity in *P. vindemmiae* and no evidence of either paternal transmission or infectious transmission. This observation contrast to *A. nasoniae* in *N. vitripennis*, which relies on both vertical and horizontal transmission to persist in the parasitoid population (17). The contrast with strain Nv makes sense in light of differences in the ecology of the host species*; Nasonia vitripennis* is a gregarious parasitoid that commonly exhibits superparasitism in which two female wasps utilize the same fly pupa, providing a opportunity for infectious transmission within the shared environment (33). *Pachycrepoideus vindemmiae*, in contrast, lays a single egg per host fly, and rarely superparasites (34). Thus, the difference in *Arsenophonus* transmission modes reflects in part distinct opportunities for horizontal transmission associated with differences in host biology.

The very close relatedness of maternally inherited *A. nasoniae* strain Pv to *A. nasoniae* Nv which has mixed modes of transmission likely therefore reflects a very recent transition of the Pv strain to a purely vertically transmitted lifestyle. The relationship is a facultative heritable symbiosis (uninfected hosts can be generated through antibiotic treatment and are viable and fertile). Segregational loss during vertical transmission means the symbiont would be lost from populations in the absence of a ‘drive’. Our data rule out one form of drive - reproductive parasitism through either sex ratio distortion or cytoplasmic incompatibility– and thus imply a balancing benefit to infection. The nature of this benefit is uncertain; offensive and defensive symbiosis are possible, but the entomophagous nature of *P. vindemmiae* perhaps makes a nutritional role less likely.

Evolution of microbial symbionts that are vertically transmitted is typically reductive, with progressive loss of metabolic function associated with systems rendered redundant by permanent host association and the lack of need for active infection (35). This process underpins the observation that the vast majority of heritable symbionts of insects are fastidious to culture. Our study highlights *A. nasoniae* Pv as a strain that is maternally inherited and very closely related to *A. nasoniae* Nv, a strain that contrastingly has mixed modes of transmission in *Nasonia* wasps. Nevertheless, strain Pv retains the capacity for growth on cell free media. Biolog analysis indicated this strain did have more restrictive requirements for culture, with a narrower range of carbon utilization sources. Thus, this datum supports the conceptual view that metabolic competence reduces over evolutionary time when infectious transmission is very rare. Onward, comparative genomic analysis of pathway loss in this strain will be instructive.

The finding of *Arsenophonus* infecting the butterfly *P. bellargus* extends the range of *A. nasoniae/A. apicola* clade symbionts beyond Hymenoptera for the first time. The infected individual was the only one carrying *Arsenophonus* in a sample of 30 male butterflies; the host died shortly after collection and had very high read depth of *Arsenophonus* redolent of those found during *A. apicola* infection. *Polyommatus bellargus* was the only eukaryotic signal present in the illumina reads, so we can be clear the *Arsenophonus* was associated with the butterfly and not a passenger parasite/commensal. *Arsenophonus apicola* exists as an environmentally acquired infection in the Hymenoptera pollinator community (14, 36), and it is tempting to suggest that *Arsenophonus* in this butterfly is likewise acquired via the pollinator environment through shared flower environments, and that it represents an opportunistic pathogen. Further experiments will be needed to assess this hypothesis. The recovery into culture of *Arsenophonus* strain Pb from this dead specimen supports a means for isolating further novel *Arsenophonus* strains through dividing hosts, testing one half by PCR assay whilst retaining the other half preserved at −80°C for onward culture in case the symbiont is detected.

Finally, our study indicates *Arsenophonus* strain Pb is capable of infecting *Galleria mellonella* waxworms, a model system for understanding host-pathogen interactions (37). This capacity contrasted with *A. nasoniae* and *Arsenophonus* strain Pv. The capacity of the *P. bellargus* strain to infect *Galleria* may reflect either the relatedness of the host species (both Lepidoptera), or a more generalist infectious transmission network of *A. nasoniae* Pb in nature. The capacity of *A. nasoniae* Pb to grow in *Galleria* provides the opportunity for understanding the molecular basis of *Arsenophonus* interaction with insects, including interface with the host immune system. Unlike many other insect endosymbionts, *A. nasoniae* and *A. apicola* are predominantly extracellular symbionts/pathogens, exposed to humoral and cellular immune systems. The *Galleria* system will enable functional analysis of how *Arsenophonus* establishes infection in the face of these host responses.

## Acknowledgements

We thank Carl Yung and Prof Ilik Saccheri for the *P. bellargus* butterfly samples and read data, and Prof Fabrice Vavre for the *Pachycrepoideus vindemmiae* strain. This work was funded by a NERC studentship (to SP) and a BBSRC grant BB/S017534/1 (to GH).

## Data availability

Data underpinning Figures 1 and 2 and onward analyses can be downloaded from https://doi.org/10.6084/m9.figshare.20628225.v1 and https://doi.org/10.6084/m9.figshare.20628798.v1

## BIBLIOGRAPHY

1. Hurst GDD. Extended genomes: symbiosis and evolution. Interface Focus. 2017;7(5):20170001.

2. Douglas AE. The microbial dimension in insect nutritional ecology. Functional Ecology. 2009;23(1):38–47.

3. Xie JL, Vilchez I, Mateos M. Spiroplasma Bacteria Enhance Survival of Drosophila hydei Attacked by the Parasitic Wasp Leptopilina heterotoma. Plos One. 2010;5(8).

4. Corbin C, Heyworth ER, Ferrari J, Hurst GDD. Heritable symbionts in a world of varying temperature. Heredity. 2017;118(1):10–20.

5. Hedges LM, Brownlie JC, O’Neill SL, Johnson KN. Wolbachia and Virus Protection in Insects. Science. 2008;322(5902):702-.

6. Teixeira L, Ferreira A, Ashburner M. The bacterial symbiont Wolbachia induces resistance to RNA viral infections in Drosophila melanogaster. Plos Biology. 2008;12:2753–63.

7. Nazni WA, Hoffmann AA, NoorAfizah A, Cheong YL, Mancini MV, Golding N, et al. Establishment of <em>Wolbachia</em> Strain <em>w</em>AlbB in Malaysian Populations of <em>Aedes aegypti</em> for Dengue Control. Current Biology. 2019;29(24):4241–8.e5.

8. Utarini A, Indriani C, Ahmad RA, Tantowijoyo W, Arguni E, Ansari MR, et al. Efficacy of Wolbachia-Infected Mosquito Deployments for the Control of Dengue. New England Journal of Medicine. 2021;384(23):2177–86.

9. Hurst GDD, Frost CL. Reproductive Parasitism: Maternally Inherited Symbionts in a Biparental World. Cold Spring Harbor Perspectives in Biology. 2015;7(5).

10. Ryder JJ, Hoare M-J, Pastok D, Bottery M, Boots M, Fenton A, et al. Disease Epidemiology in Arthropods Is Altered by the Presence of Nonprotective Symbionts. American Naturalist. 2014;183(3):E89–E104.

11. Charlat S, Reuter M, Dyson EA, Hornett EA, Duplouy A, Davies N, et al. Male-killing bacteria trigger a cycle of increasing male fatigue and female promiscuity. Current Biology. 2007;17(3):273–7.

12. Miller AK, Westlake CS, Cross KL, Leigh BA, Bordenstein SR. The microbiome impacts host hybridization and speciation. PLOS Biology. 2021;19(10):e3001417.

13. Sachs JL, Skophammer RG, Regus JU. Evolutionary transitions in bacterial symbiosis. Proceedings of the National Academy of Sciences of the United States of America. 2011;108(Suppl 2):10800–7.

14. Drew GC, Budge GE, Frost CL, Neumann P, Siozios S, Yañez O, et al. Transitions in symbiosis: evidence for environmental acquisition and social transmission within a clade of heritable symbionts. The ISME Journal. 2021;15:2956–68.

15. Nadal-Jimenez P, Siozios S, Frost CL, Court R, Chrostek E, Drew GC, et al. Arsenophonus apicola sp. nov., isolated from the honeybee Apis mellifera. International Journal of Systematic and Evolutionary Microbiology. 2022;72(8).

16. Bressan A, Moral García FJ, Boudon-Padieu E. The prevalence of ‘Candidatus Arsenophonus phytopathogenicus’ infecting the planthopper Pentastiridius leporinus (Hemiptera: Cixiidae) increase nonlinearly with the population abundance in sugar beet fields. Environ Entomol. 2011;40(6):1345–52.

17. Parratt SR, Frost CL, Schenkel MA, Rice A, Hurst GDD, King KC. Superparasitism Drives Heritable Symbiont Epidemiology and Host Sex Ratio in a Wasp. Plos Pathogens. 2016;12(6).

18. Hypsa V, Dale C. In vitro culture and phylogenetic analysis of ‘‘Candidatus *Arsenophonus triatominarum,’’* an intracellular bacterium from the triatomine bug, *Triatoma infestans*. Intl J Syst Bact. 1997;47:1140–4.

19. Dale C, Beeton M, Harbison C, Jones T, Pontes M. Isolation, pure culture, and characterization of “Candidatus Arsenophonus arthropodicus,” an intracellular secondary endosymbiont from the hippoboscid louse fly Pseudolynchia canariensis. Applied And Environmental Microbiology. 2006;72(4):2997–3004.

20. Nováková E, Hypša V, Nguyen P, Husník F, Darby AC. Genome sequence of Candidatus Arsenophonus lipopteni, the exclusive symbiont of a blood sucking fly Lipoptena cervi (Diptera: Hippoboscidae). Stand Genomic Sci. 2016;11:72.

21. Nadal-Jimenez P, Griffin JS, Davies L, Frost CL, Marcello M, Hurst GDD. Genetic manipulation allows in vivo tracking of the life cycle of the son-killer symbiont, Arsenophonus nasoniae, and reveals patterns of host invasion, tropism and pathology. Environ Microbiol. 2019;21(8):3172–82.

22. Frost CL, Siozios S, Nadal-Jimenez P, Brockhurst MA, King KC, Darby AC, et al. The Hypercomplex Genome of an Insect Reproductive Parasite Highlights the Importance of Lateral Gene Transfer in Symbiont Biology. mBio. 2020;11(2).

23. Duron O, Wilkes TE, Hurst GDD. Interspecific transmission of a male-killing bacterium on an ecological timescale. Ecology Letters. 2010;13(9):1139–48.

24. Darby AC, Choi JH, Wilkes T, Hughes MA, Werren JH, Hurst GDD, et al. Characteristics of the genome of Arsenophonus nasoniae, son-killer bacterium of the wasp Nasonia. Insect molecular biology. 2010;19:75–89.

25. Turner S, Pryer KM, Miao VP, Palmer JD. Investigating deep phylogenetic relationships among cyanobacteria and plastids by small subunit rRNA sequence analysis. J Eukaryot Microbiol. 1999;46(4):327–38.

26. Duron O, Bouchon D, Boutin S, Bellamy L, Zhou LQ, Engelstadter J, et al. The diversity of reproductive parasites among arthropods: Wolbachia do not walk alone. BMC Biol. 2008;6.

27. Folmer O, Black M, Hoeh W, Lutz R, Vrijenhoek R. DNA primers for amplification of mitochondrial cytochrome c oxidase subunit I from diverse metazoan invertebrates. Molecular Marine Biology and Biotechnology. 1994;3(5):294–9.

28. Gruber-Vodicka HR, Seah BKB, Pruesse E. phyloFlash: Rapid Small-Subunit rRNA Profiling and Targeted Assembly from Metagenomes. mSystems. 2020;5(5).

29. Kalyaanamoorthy S, Minh BQ, Wong TKF, von Haeseler A, Jermiin LS. ModelFinder: fast model selection for accurate phylogenetic estimates. Nature Methods. 2017;14(6):587–9.

30. Nguyen L-T, Schmidt HA, von Haeseler A, Minh BQ. IQ-TREE: A Fast and Effective Stochastic Algorithm for Estimating Maximum-Likelihood Phylogenies. Molecular Biology and Evolution. 2015;32(1):268–74.

31. Hoang DT, Chernomor O, von Haeseler A, Minh BQ, Vinh LS. UFBoot2: Improving the Ultrafast Bootstrap Approximation. Molecular Biology and Evolution. 2017;35(2):518–22.

32. Vavre F, Fleury F, Varaldi J, Fouillet P, Boulétreau M. Evidence for female mortality in *Wolbachia-*mediated cytoplasmic incompatibility in haplodiploid insects: epidemiologic and evolutionary consequences. Evolution. 2000;54:191–200.

33. Grillenberger B, Van de Zande L, Bijlsma R, Gadau J, Beukeboom L. Reproductive strategies under multiparasitism in natural populations of the parasitoid wasp <i>Nasonia</i> (Hymenoptera). Journal of Evolutionary Biology. 2009;22(3):460–70.

34. Li J, Gong X-M, Chen Y-Z, Pan S-Y, Dai Y-N, Hu H-Y, et al. Effect of maternal age on primary and secondary sex ratios in the ectoparasitoid wasp Pachycrepoideus vindemmiae. Entomologia Experimental et Applicata. 2022;170(6):468–76.

35. McCutcheon JP, Moran NA. Extreme genome reduction in symbiotic bacteria. Nature Reviews Microbiology. 2012;10(1):13–26.

36. Yanez O, Gauthier L, Chantawannakul P, Neumann P. Endosymbiotic bacteria in honey bees: Arsenophonus spp. are not transmitted transovarially. Fems Microbiology Letters. 2016;363(14).

37. Kemp MW, Massey RC. The use of insect models to study human pathogens. Drug Discovery Today: Disease Models. 2007;4(3):105–10.

